# Neighborhood Deprivation, Genetic Predisposition, and Life Satisfaction: Evidence from the German Twin Family Panel

**DOI:** 10.1101/2024.12.12.628202

**Authors:** Nadia V. Harerimana, Yixuan Liu, Mirko Ruks

## Abstract

Both genes and the neighborhood are important for life satisfaction; however, there is little research on gene-environment interactions (GxE) that examines how the effect of genetic endowments varies as a function of the environmental context with life satisfaction as the outcome. This study investigated how neighborhood deprivation moderates the effects of genetic predisposition on life satisfaction. Using data from the German Twin Family Panel (TwinLife), we identified 760 dizygotic (DZ) twins and employed twin fixed-effect models to assess the GxE effects on life satisfaction. The findings reveal that the polygenic score (PGS) for subjective well-being is positively associated with life satisfaction. The effect of PGS for subjective well-being on life satisfaction is strongest for individuals living in moderately deprived areas, while it is weaker for those living in highly deprived and less deprived areas. Thus, there are signs of compensation in less deprived areas and, particularly, diathesis-stress/triggering in highly deprived areas.

## 1. Introduction

Life satisfaction, a fundamental element of subjective well-being, encapsulates how individuals cognitively evaluate their lives. Beyond its intrinsic value, research highlights the multifaceted benefits associated with life satisfaction. Studies indicate its correlation with various positive life outcomes, including longevity, enhanced health, academic achievement, income levels, and involvement in civic activities (Antaramian, 2017; Borman et al., 2001; Collins et al., 2009; Danner et al., 2001; De Neve and Oswald, 2012; Diener and Chan, 2011; Diener et al., 2018a,b, 2017; Flavin and Keane, 2012; Frey, 2011; Lyubomirsky et al., 2005). Therefore, it is important to study the factors influencing life satisfaction.

One crucial factor explaining differences in life satisfaction is genetics as behavioral genetics research has consistently shown that genetic differences can be linked to differences in life satisfaction. Different metaanalyses report heritability estimates ranging from 32% (Bartels, 2015) to 40% (Nes and Røysamb, 2015), meaning that up to 40% of the differences in life satisfaction can be traced back to genetic differences.

Different studies suggest that a substantial portion of this genetic influence is accounted for by genetic variants related to personality traits (Lachmann et al., 2021; Røysamb et al., 2018). On the other hand, there are environmental factors influencing life satisfaction. In this regard, several studies have focused on neighborhood effects on life satisfaction. Research has consistently shown that residing in disadvantaged neighborhoods increases the risk of mortality and mental illnesses such as depression and anxiety (Kim, 2008; Mair et al., 2008; Truong and Ma, 2006). This association may stem from various factors including poor physical conditions, limited access to resources, exposure to environmental toxins, and a lack of social cohesion and safety (Carrillo- Álvarez et al., 2019; Foster and Giles-Corti, 2008; Mohammed et al., 2019; Quinn et al., 2019). Unfavorable neighborhood conditions have been linked to lower life satisfaction and overall poorer health outcomes among residents (Bagheri et al., 2019; Orben et al., 2020). Comparative studies conducted in England, Wales, and Sweden have all demonstrated a negative correlation between neighborhood deprivation and life satisfaction, with individuals in better-off neighborhoods generally experiencing better health outcomes (Knies et al., 2021; White et al., 2016). Similar trends have been observed in Germany, where studies have revealed that individuals living in neighborhoods with higher socioeconomic status report higher levels of life satisfaction, even after accounting for household and individual characteristics (Dittmann and Goebel, 2010).

So, genes and the neighborhood are important for life satisfaction. However, both influences do not need to be independent from each other. In this regard, the research on gene-environment interactions (GxE) studies how the effect of genetic endowments varies as a function of the environmental context and vice versa (Shanahan and Hofer, 2005). In fact, multiple studies have investigated whether genetic effects vary as a function of neighborhood deprivation. Most of these studies focus on health or psychopathological outcomes or overall problem behavior. A twin study by Singh et al. (2021) reports that perceived neighborhood cohesion moderates the genetic effect on internalization but not on externalizing behavior. For females, the relevance of genetics decreases with higher neighborhood cohesion, they find an opposite pattern for male. Strachan et al. (2017) found that genetic influences on depression increases with increasing neighborhood deprivation. In line with that, Carroll et al. (2023) find that genetic influences on ADHD are higher among individuals living in disadvantaged neighborhoods. Rhew et al. (2020) report that while neighborhood deprivation moderates environmental influences on hazardous drinking, it does not moderate genetic influences. Dash et al. (2023) show that genetic influences on externalizing problem behavior are weaker for individuals living in neighborhoods with lower educational, or social and economic opportunities. Burt et al. (2020) show that genetic influences on antisocial behavior in childhood are greater among individuals living in advantaged neighborhoods. While the studies discussed so far, applied twin modeling to estimate genetic effects, Pasman et al. (2020) use a polygenic score (PGS) to measure the genetic disposition for substance use. Studying 14 GxE models, they found only the genetic effect on alcohol use was significantly moderated by neighborhood socio-economic status with stronger genetic effects for individuals living in higher status neighborhoods.

Importantly, none of these studies focus on life satisfaction as the outcome. However, as the meta-analysis by Roysamb et al. (2018) shows, there is significant heterogeneity in the heritability estimates for life satisfaction suggesting the presence of gene-environment interactions (GxE). Against this background, the study of GxE has remained surprisingly scarce for life satisfaction compared to other complex traits (Bartels, 2015) with studies testing whether heritability of life satisfaction varies with family socio- eocnomic status (Johnson and Krueger, 2006), marital status (Nes et al., 2010) or parental divorce (van der Aa et al., 2010).

Therefore, this study aims to address this gap in the literature by investigating whether the genetic influence on life satisfaction varies as a function of neighborhood deprivation. This study leverages a co-twin control design to investigate GxE effects on life satisfaction, effectively addressing prior methodological limitations by using the natural experiment of sibling differences in genetic dispositions to account for genetic confounding effects. Further, this study investigates how contextual factors at the level of neighborhood moderate the effects of genetic predispositions on life satisfaction by combining geo-coded data on neighborhood deprivation with individual-level and household-level survey data from the longitudinal twin family study of German twins and their immediate relatives. This study represents a pioneering effort to reveal the interplay between genetic predispositions and environmental factors at the neighborhood level on life satisfaction at the individual level in Germany.

## 2. Theoretical Framework

In this section, we discuss theoretical arguments on how and why the genetic effect on life satisfaction may vary as a function of neighborhood deprivation. To do so, in a first step we distinguish general GxE mechanisms. Then, we link these GxE mechanisms to the mechanisms of neighborhood effects discussed in the literature to explain how neighborhood deprivation might affect the genetic effect on life satisfaction.

### 2.1. GxE Mechanisms

Shanahan and Hofer (2005) distinguish four GxE mechanisms. Triggering describes situations where exposure to disadvantaged environmental contexts, characterized by high levels of risk and stressors, increases the probability of realizing genetic vulnerabilities. Compensation refers to cases where the realization of a genetic risk is limited in advantaged environments. Social control is a specific form of compensation that focuses on the enforcement of social norms and values in advantaged environments restricting the realization a genetic risk. Finally, enhancement refers to a multiplication process where the realization of a positive genetic endowment is amplified in advantaged environments.

It’s possible to categorize these mechanisms based on the type of genetic disposition and environmental context. However, Shanahan and Hofer (2005) don’t delve into how a disadvantaged environment can affect the realization of a positive genetic potential. Therefore, we introduce the concept of suppression, according to which the expression of a genetic endowment is limited in disadvantaged environments. Overall, while triggering and compensation imply stronger genetic effects in disadvantaged environments (negative GxE), suppression and enhancement explain stronger genetic effects in advantaged environments (positive GxE).

An overview of these GxE mechanisms is provided in Table 1.

**Table 1:**
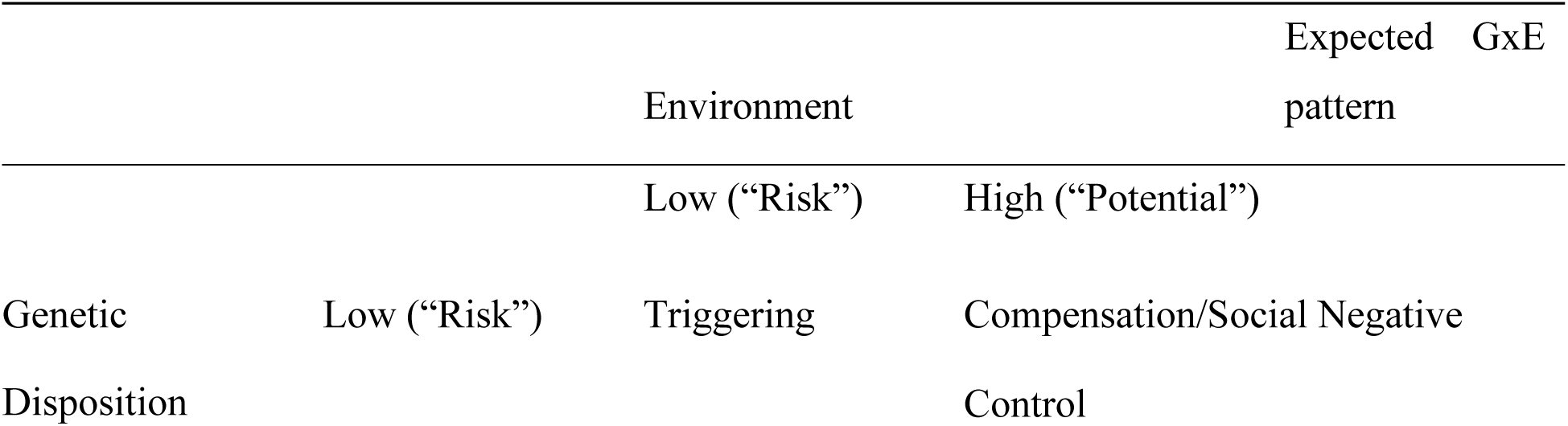
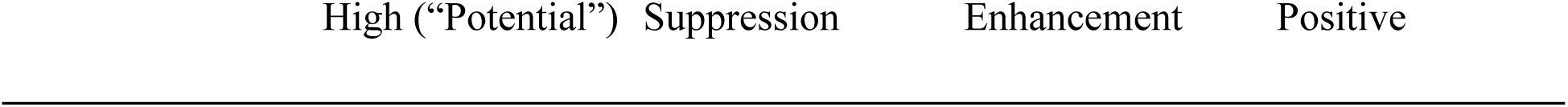
Expected GxE mechanisms.

|  |  | Environment |  | Expected GxE pattern |
| --- | --- | --- | --- | --- |
|  |  | Low ("Risk") | High ("Potential") |  |
| Genetic Disposition | Low ("Risk") | Triggering | Compensation/Social Control | Negative |

### 2.2. GxNeighborhood mechanisms

Various theoretical mechanisms driving neighborhood effects have been discussed in the literature (for an overview, see e.g., Ellen and Turner, 1997; Galster, 2012; Jencks and Mayer, 1990; Musterd et al., 2019; Silva, 2023). Here, we first discuss five mechanisms underlying the effects of neighborhood deprivation. Then we link these mechanisms to the overall GxE typology derived before to explore how neighborhood deprivation might moderate the genetic effect on life satisfaction.

First, epidemic models emphasize the role of peers within the neighborhood for socialization. As human behavior is shaped partly by social learning (Akers et al., 1979), it is assumed that neighborhoods can impact individual behavior through interaction with peers in the area. In this view, individuals tend to mirror the behavior of their peers in terms of a contagion effect (“like begets like”) (Jencks and Mayer, 1990). In disadvantaged neighborhoods, peers may exhibit higher levels of problem behavior (Wilson, 1987), potentially leading to negative outcomes such as increased victimization (Foster and Brooks-Gunn, 2013) or depressive symptoms (Homel et al., 2020). Conversely, in advantaged neighborhoods, individuals are more likely to encounter successful and prosocial peers, fostering social support and opportunities for positive friendships (Brown et al., 2008; Fredricks and Simpkins, 2013).

Second, collective socialization models highlight the influence of non-parent adults in a neighborhood on children and adolescents (Jencks and Mayer, 1990). These adults can serve as role models, shaping aspirations and defining what behavior is considered acceptable (Ellen and Turner, 1997; Jencks and Mayer, 1990). In disadvantaged neighborhoods, exposure to negative role models may be more prevalent, promoting attitudes that devalue education and encourage problematic behavior. Additionally, limited job opportunities for adults in these areas may lead adolescents to perceive little benefit from responsible behavior, resulting in lower commitment to education (Ellen and Turner, 1997; Anderson, 2000). Conversely, in affluent neighborhoods, positive adult role models can inspire, guide, and support individuals, fostering higher aspirations, self-efficacy, and motivation. These adults may also serve as enforcers, monitoring behavior, maintaining order, and intervening when necessary.

A third mechanism operates through the institutional resources available in a neighborhood, which can significantly impact adolescent development (Sampson et al., 2002; Sirin, 2005). These resources, such as schools, libraries, and social services, vary in quality, quantity, and diversity depending on the neighborhood’s status. In affluent neighborhoods, schools often have better facilities and teachers, and there is a wider range of social and cultural activities and extracurricular opportunities, such as sports, music, or art. Conversely, in disadvantaged neighborhoods, these institutional resources may be lacking, placing adolescents at greater risk of engaging in problematic or delinquent behavior and limiting their ability to develop talents and skills essential for future socioeconomic success (Ellen and Turner, 1997).

The fourth mechanism highlights the significance of social capital, which encompasses the connections and networks among individuals, along with the norms of reciprocity and trust (Putnam, 1995). Neighborhood social capital encompasses various dimensions, including the level of social ties, frequency of interactions, and neighboring patterns (Bellair, 1997, 2000; Rountree and Warner, 1999). These social ties within a neighborhood can be advantageous, fostering a cohesive community where residents support each other and share valuable information and resources. In affluent neighborhoods, there tends to be a higher quantity and quality of social ties, along with shared resources, providing residents with a stronger sense of community and access to valuable knowledge about status attainment and job opportunities. Conversely, in deprived neighborhoods, social ties and shared resources may be lacking or significantly reduced.

Lastly, collective efficacy refers to a neighborhood’s ability to mobilize informal social control, organize, and pursue common goals, relying on mutual trust, shared norms, and expectations (Sampson et al., 1997). Neighborhoods with clear rules and trust among residents are more likely to successfully activate social control, leading to positive outcomes such as reduced violence, disorder, and better access to public services. Additionally, residents may feel more empowered and engaged in local events, fostering a sense of belonging. Studies have shown that higher collective efficacy is associated with reduced criminal victimization, deviant behavior, and better mental health (Browning et al., 2014; Drukker et al., 2003; Dupéré et al., 2012; Jain et al., 2010; Sampson et al., 1997; Simons et al., 2005; Xue et al., 2005).

These mechanisms can be related to the overall GxE mechanisms discussed before, explaining why neighborhood deprivation might moderate the genetic effect on life satisfaction. On the one hand, in affluent neighborhoods the exposure to 1) positive peer influences, 2) motivating and involved role models, 3) the high quality and availability of supporting institutions, 4) higher quality and quantity of social capital, and 5) higher levels of collective efficacy may have a compensating function so that individuals develop high levels of life satisfaction despite low genetic endowment for life satisfaction. In line with that, in deprived neighborhoods, the lack of these experiences may act as a stressor, triggering the manifestation of genetic disadvantages in terms of low life satisfaction. As a result, we would expect the genetic effect size to increase with higher levels of neighborhood deprivation (H1). On the other hand, the positive neighborhood experiences in affluent neighborhoods may actually enhance the realization of a genetic potential for life satisfaction so that individuals in affluent neighborhoods profit disproportionately from a high genetic disposition for life satisfaction. In deprived neighborhoods, in turn, the lack of these experiences may act as a suppressor, preventing the realization of an actually high genetic endowment for life satisfaction. From this, we would expect the genetic effect size to decrease with higher levels of neighborhood deprivation (H2).

## 3. Data and methods

### 3.1. Participants

The German Twin Family Panel (TwinLife), which began in 2014, is a nationally representative cohort sequential panel-study of four cohorts of same-sex twins and their biological families (Diewald et al., 2023; Mönkediek et al., 2019). Unlike many convenience twin studies based on self-selection of twins, TwinLife applies a register-based probability design. It has been shown that the TwinLife includes families from the whole socio-economic spectrum in Germany (Lang and Kottwitz, 2020) and the socio-demographic structure is comparable to other German family surveys (Mönkediek et al., 2020). This study uses data of cohorts 2-4 from the first F2F wave conducted in 2014-2016.

### 3.2 Ethical statement

The data analyzed in this study were obtained from participants of the TwinLife project. Genetic data, including saliva samples for DNA extraction, were collected as part of two TwinLife satellite projects: TwinSNPs and TECS. Ethical approval for the TwinLife study was granted by the German Psychological Society (Protocol Number: RR 11.2009). The protocols for genetic sampling via saliva were additionally reviewed and approved by the Ethics Committee of the Medical Faculty at the University of Bonn (Approval Number: 113/18). Data collection for the Face-to-Face (F2F 1) survey occurred in two phases: Subsample A was collected between September 28, 2014, and May 28, 2015, while Subsample B was collected between September 16, 2015, and April 18, 2016

### 3.3. Measures

#### 3.3.1. Life satisfaction

Life satisfaction was measured via the “Satisfaction with Life Scale” (SWLS) (Diener et al., 1985) for participants aged 16 years and above as well as using an adapted version for children aged 10 - 15 years (SWLS-C) (Gadermann et al., 2010), comprising five items each with a five-point Likert scale (1 = Disagree strongly - 5 = Agree strongly). The final life satisfaction score was created using confirmatory factor analysis of the raw indicators.

#### 3.3.2. Genetic predisposition for life satisfaction

The genetic predisposition for life satisfaction was measured via a polygenic score (PGS) for subjective well-being. Participants’ saliva was extracted between 2018 and 2020 (Diewald et al., 2023; Mönkediek et al., 2019). PGS for subjective well-being was estimated for TwinLife participants in Wave 1 using results from the well-being spectrum (Becker et al., 2021) following the PRS-CS approach (Ge et al., 2019). Briefly, genetic variants were filtered for (i) an imputation info score >= 0.6 if available in the respective summary statistics, (ii) a minor allele frequency >= 0.01 in the 1000 Genomes phase 3 European LD reference panel, and (iii) presence in both the respective summary statistics and the imputed genotype data of TwinLife. For a detailed description of sampling, genotyping, quality control, filtering, and PGS calculation, please see Supplement B-I of the study (Deppe et al., 2024). Subsequently, associations between PGS and neighborhood index were examined, adjusting for sex, 10 genetic principal components using regression modeling.

#### 3.3.3. Neighborhood deprivation

Neighborhood deprivation is measured using the German Index of Socioeconomic Deprivation (GSID) developed by Michalski et al. (2022) which is publicly available (Michalski et al., 2024). The index was created based on administrative data for Germany published by the INKAR database and comprises three dimensions (Education, Employment, and Income) each of which is measured via three indicators. The GSID was extracted via principal component analysis and matched to the TwinLife survey data based on district codes. To account for possible non- linearities, we categorized the GSID score along the terciles, distinguishing low, middle, and high neighborhood deprivation.

#### 3.3.4. Covariates

As families might select into neighborhood deprivation partly as a function of family socio- economic status or age structure, in the statistical analysis we additionally control for family SES, measured as a factor based on parental years of education, occupational status (ISEI) and OECD household net income. Additionally, we control for birth cohort status, using a categorical variable indicating whether a twin belongs to cohort 2, 3, or 4.

Finally, Table 2 provides a descriptive overview of the full sample (N=1544) and Table 3 splits up the descriptive by zygosity (N DZ twins = 760; N MZ twins = 784) showing no meaningful differences across zygosity.

**Table 2:** Descriptive statistics of the full sample (N=1544)

|  | Min | Max | Mean | SD |
| --- | --- | --- | --- | --- |
| Life Satisfaction | -4.49 | 1.38 | 0.00 | 1.00 |
| PGS | -4.13 | 3.15 | 0.00 | 1.00 |
| Deprivation Index | -2.51 | 2.30 | 0.00 | 1.00 |
| SES | -2.41 | 2.51 | 0.00 | 1.00 |
| Cohort 2 (10-12) | 0.00 | 1.00 | 0.46 | 0.50 |
| Cohort 3 (16-18) | 0.00 | 1.00 | 0.36 | 0.48 |
| Cohort 4 (22-25) | 0.00 | 1.00 | 0.18 | 0.38 |
| Female | 0.00 | 1.00 | 0.56 | 0.50 |
| Age | 10.00 | 25.00 | 15.31 | 4.52 |

**Table 3:** Descriptive statistics by zygosity (N DZ twins = 760; N MZ twins = 784)

|  | DZ |  |  |  | MZ |  |  |  |
| --- | --- | --- | --- | --- | --- | --- | --- | --- |
|  | Min | Max | Mean | SD | Min | Max | Mean | SD |
| Life Satisfaction | -4.14 | 1.38 | -0.03 | 1.00 | -4.49 | 1.38 | 0.03 | 0.99 |
| PGS | -3.03 | 3.15 | 0.05 | 0.99 | -4.13 | 3.02 | -0.05 | 1.00 |
| Deprivation Index | -2.51 | 2.26 | 0.00 | 0.95 | -2.51 | 2.30 | -0.01 | 1.05 |
| SES | -2.40 | 2.33 | 0.07 | 0.99 | -2.41 | 2.51 | -0.07 | 1.00 |
| Cohort 2 (10-12) | 0.00 | 1.00 | 0.53 | 0.50 | 0.00 | 1.00 | 0.40 | 0.49 |
| Cohort 3 (16-18) | 0.00 | 1.00 | 0.37 | 0.48 | 0.00 | 1.00 | 0.35 | 0.48 |
| Cohort 4 (22-25) | 0.00 | 1.00 | 0.11 | 0.31 | 0.00 | 1.00 | 0.25 | 0.43 |
| Female | 0.00 | 1.00 | 0.54 | 0.50 | 0.00 | 1.00 | 0.57 | 0.49 |
| Age | 10.00 | 24.00 | 14.46 | 4.03 | 10.00 | 25.00 | 16.13 | 4.82 |

### 3.4. Statistical Analysis

To test the competing GxE hypotheses, we apply a step-wise regression approach. In a first step, we estimate a multi-level model accounting for the family structure on the pooled sample. In a second step, we estimate a fixed effects model on the DZ twin sample, accounting for unobserved between-family heterogeneity. Overall, with twin i clustered in twin pair j, the model equation is given by:

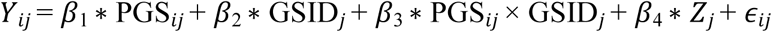

Here, PGS_*ij*_ is the measured genotype, GSID_*j*_ is the neighborhood deprivation, varying only between twin pairs, and Z_*j*_ is a vector of controls. Importantly, in order to get unbiased estimates of the PGS for subjective well-being (*β*_1_) and of its moderation by neighborhood deprivation (*β*_3_), the exogeneity assumption must hold. In other words, the estimates are biased if Cov(PGS_*ij*_,*ϵ*_*ij*_) = Cov(PGS_*ij*_ × GSID_*j*_,*ϵ*_*ij*_) ≠ 0.

Family data can be leveraged to address this problem as it is possible to decompose the error term: *ϵ*_*ij*_= *u*_*j*_+*e*_*ij*_, where *u*_*j*_is an error term varying between DZ twin pairs whereas *e*_*ij*_denotes an idiosyncratic error term, varying within DZ twin pairs. When demeaning the variables in a fixed effects model (*Y*_*ij*_ −*Y*_*j*_= *Y*_*ij*_), *u*_*j*_ drops from the equation, meaning that the exogeneity assumption with respect to *u*_*j*_ can be relaxed. Thus, for the estimate of PGS_*ij*_ ×GSID_*j*_ to be unbiased, one has only to assume exogeneity with respect to *e*_*ij*_.

In addition, it is also often argued that in a within-family design the PGS is quasi-exogenous as genetic differences in full biological siblings are randomized at conception as the transmission of alleles from parents to children is a random process (Domingue et al., 2015; Fletcher and Lehrer, 2011; Schmitz and Conley, 2017). So, there are strong reasons to assume PGS as exogenous.

Given that families might select into neighborhood deprivation based on family SES or birth cohort, we additionally control for PGS_*ij*_×SES_*j*_ and PGS_*ij*_×Cohort_*j*_. FE models draw only on within-family variation. Therefore, the main effects of variables varying only between families (GSID, family SES, and cohort) are not estimated.

The model equation for the fixed effects model is:

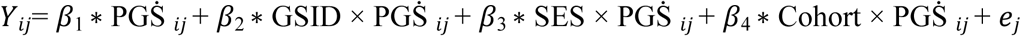

We account for the different controls in a stepwise procedure, resulting in four models for the pooled and FE analysis: In the first model (M1), the raw PGS effect is estimated. The second model (M2) adds the interaction between PGS and GISD. The third model (M3) additionally accounts for the interaction between PGS and family SES. The fourth model (M4) additionally controls for the interaction between PGS and birth cohort.

## 4. Results

### 4.1. Pooled analysis

The results of the pooled analysis is shown in Table 4. The first model suggests - on average - a small, but significant (p<0.05) positive genetic effect on life satisfaction on phenotypic life satisfaction. The second model accounts for possible differences in the genetic effect across levels of neighborhood deprivation. M2 shows that for individuals living in neighborhoods with a medium level of neighborhood deprivation, the genetic effect on life satisfaction is about 0.154 (p<0.01), while it is significantly lower for individuals living in less deprived neighborhoods (- 0.183, p<0.01). The genetic effect for individuals living in highly deprived neighborhoods is also lower compared to individuals living in mid-deprived neighborhoods. However, the difference is not significant (-0.076, p>0.05). The observed differences in the genetic effect may be driven by family SES or twins’ birth cohort. Therefore, M3 additionally controls for SES differences in the genetic effect and M4 accounts for cohort differences in the genetic effect. However, this does not substantially affect the estimates of the deprivation differences in the genetic effect on life satisfaction. Figure 1 shows the estimated genetic effects on life satisfaction for the three levels of neighborhood deprivation, suggesting an inverted U-shaped pattern with genetic effects being strongest for individuals living in middle deprived neighborhoods, suggesting the in highly deprived neighborhoods, the realization of a genetic potential is suppressed, while affluent neighborhoods compensate for a genetic risk. While the pooled analysis offers some first insights, the estimates might be biased by unobserved between-family characteristics.

**Figure 1:**
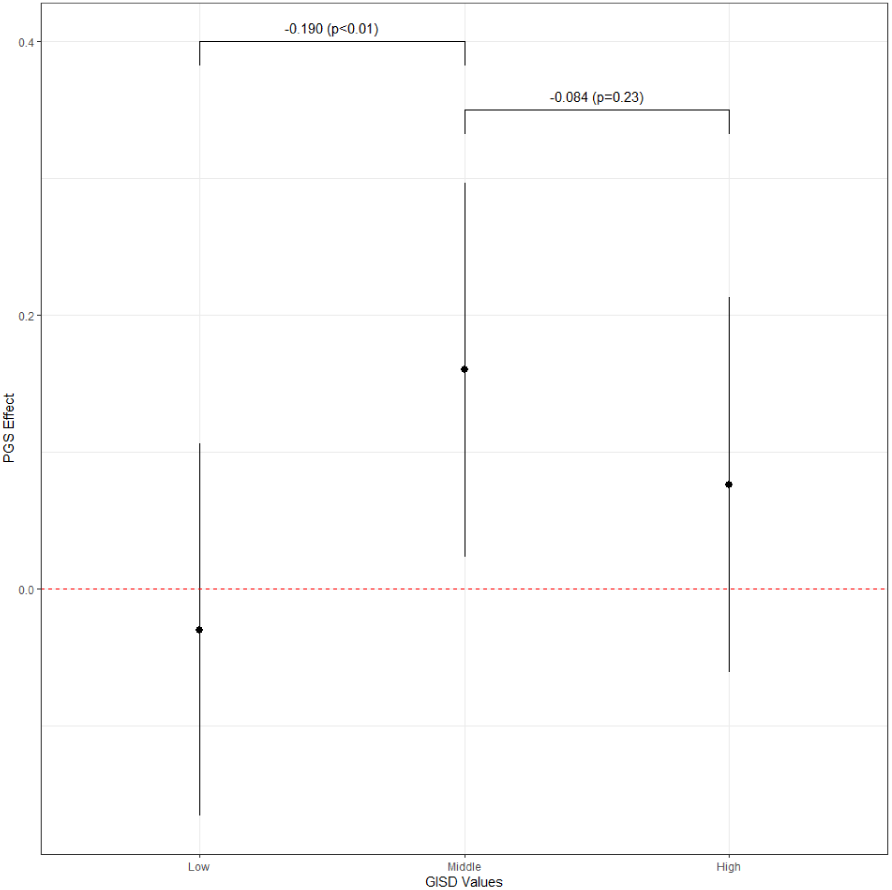
PGS efffect by GISD groups in the pooled model (M4)

**Table 4:** Results of the pooled models (N = 1544)

|  | M1 | M2 | M3 | M4 |
| --- | --- | --- | --- | --- |
| PGS | 0.063* | 0.154** | 0.125* | 0.160* |
|  | (0.028) | (0.050) | (0.063) | (0.070) |
| PGS x Low GISD |  | -0.183** | -0.189** | -0.190** |
|  |  | (0.068) | (0.069) | (0.069) |
| PGS x High GISD |  | -0.076 | -0.086 | -0.084 |
|  |  | (0.070) | (0.070) | (0.071) |
| Control for PGS x SES | No | No | Yes | Yes |
| Control for PGS x Cohort | No | No | No | Yes |
| + p < 0.1, * p < 0.05, ** p < 0.01, *** p < 0.001 |  |  |  |  |

### 4.2. Fixed effects models

Therefore, in a second step, fixed effects models are estimated to account for unobserved between-family characteristic effects (see Table 5). The first fixed effects model shows that the observed positive genetic effect on life satisfaction in the first pooled model attenuates and loses its statistical significance after adjusting for unobserved between-family characteristics. This suggests that unobserved between-family characteristics do confound the PGS-life satisfaction association. M2 shows that, for DZ twins living in neighborhoods with a medium level of neighborhood deprivation, the within-family genetic effect on life satisfaction is about 0.255 (p<0.05), while it is significantly lower for DZ twins living in less deprived neighborhoods (-0.333, p<0.05) and significantly lower for DZ twins living in more deprived neighborhoods (-0.342, p<0.05). Compared to M2, M3 additionally controls for SES differences in the within-family genetic effect, and M4 additionally accounts for both SES and cohort differences in the within-family genetic effect. For DZ twins living in neighborhoods with a medium level of neighborhood deprivation, the within-family genetic effect on life satisfaction increases to 0.404 (p<0.01) in M3, and to 0.542 (p<0.001) in M4. Like the pooled model results, including SES and cohort differences in the within-family genetic effect does not substantially affect the estimates of the deprivation differences in the within-family genetic effect on life satisfaction. Figure 2 shows the estimated within-family genetic effects on life satisfaction for the three levels of neighborhood deprivation, and the estimates are not driven by unobserved between-family characteristics. Similar to the pooled model results, this suggests a robust inverted U-shaped pattern, with genetic effects being strongest for individuals living in moderately deprived neighborhoods. This indicates that in highly deprived neighborhoods, the realization of genetic potential is suppressed, while affluent neighborhoods compensate for genetic risk.

**Figure 2:**
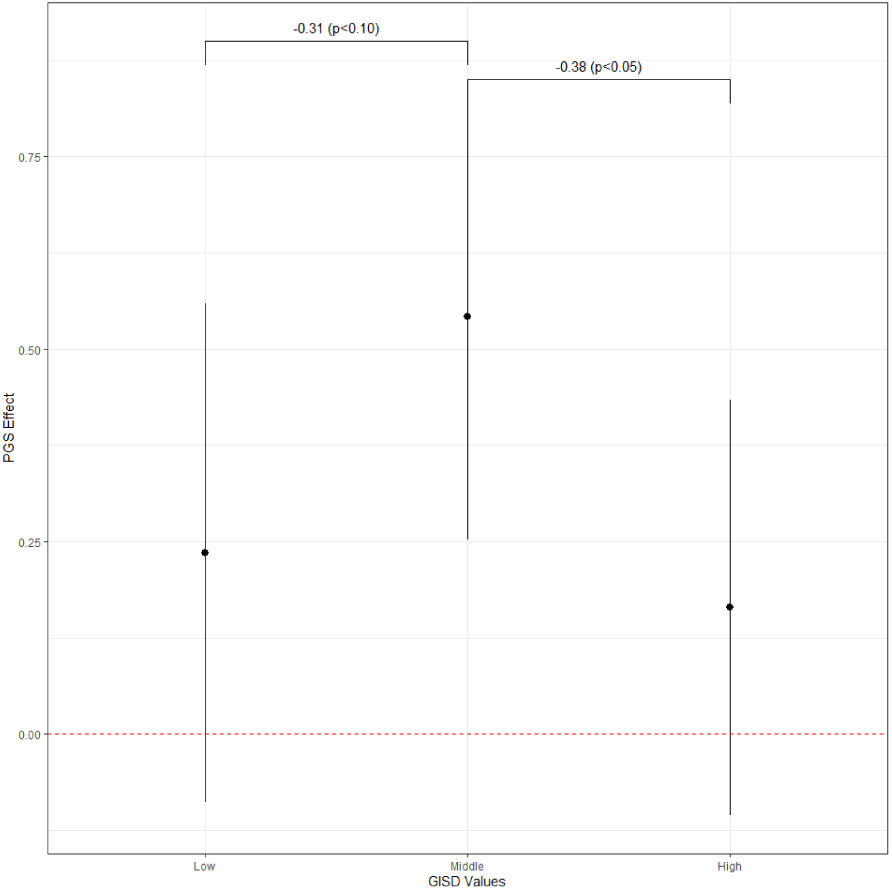
PGS effect by GISD groups in the fixed-effects model (M4)

**Table 5:** Results of fixed effects models among DZ twins (N = 760)

|  | M1 | M2 | M3 | M4 |
| --- | --- | --- | --- | --- |
| PGS (within) | 0.035<br>(0.062) | 0.255*<br>(0.105) | 0.404**<br>(0.135) | 0.542***<br>(0.148) |
| PGS (within) x Low GISD |  | -0.333*<br>(0.157) | -0.312*<br>(0.158) | -0.307+<br>(0.158) |
| PGS (within) x High GISD |  | -0.342* | -0.362* | -0.378* |
|  |  | (0.147) | (0.148) | (0.147) |
| Control for PGS x SES | No | No | Yes | Yes |
| Control for PGS x Cohort | No | No | No | Yes |

## 5. Discussion

This study examines the complex interplay between genetic predispositions and neighborhood conditions in determining life satisfaction among German individuals. Our analysis leverages the TwinLife dataset, utilizing a polygenic score (PGS) for subjective well-being and neighborhood deprivation data to investigate gene-environment interactions (GxE).

Our findings align with previous research indicating a significant genetic component to life satisfaction, with heritability estimates suggesting that up to 40% of the variance in life satisfaction can be attributed to genetic differences (Bartels, 2015; Nes and Røysamb, 2015). Our results contribute to this body of literature by confirming a positive genetic effect on life satisfaction across our sample.

The primary contribution of this study is the exploration of GxE interactions in life satisfaction, an area less studied compared to other traits. We tested two hypotheses on how neighborhood deprivation might influence the genetic impact on life satisfaction: (1) Affluent neighborhoods might compensate for genetic risks, resulting in stronger genetic effects in deprived environments; (2) Affluent neighborhoods might enhance genetic potentials, leading to stronger genetic effects in advantaged environments.

Our findings suggest a reverse relationship in the genetic effects on life satisfaction across different levels of neighborhood deprivation. Specifically, the genetic influence on life satisfaction is strongest in moderately deprived neighborhoods and weaker in both highly deprived and less deprived neighborhoods. This pattern suggests that in highly deprived neighborhoods, the realization of genetic potential for life satisfaction is suppressed, possibly due to the overwhelming negative environmental factors. In less deprived neighborhoods, positive environmental conditions might compensate for genetic risks, thereby reducing the observable genetic influence.

We employed a co-twin control design to mitigate potential confounding effects and ensure robust estimates of GxE interactions. This approach leverages the natural experiment of sibling differences in genetic dispositions, providing a powerful means to account for genetic confounding. Additionally, our use of geo-coded neighborhood deprivation data from the German Index of Socioeconomic Deprivation (GSID) adds precision to our measurement of environmental context.

Despite the strengths of our study, several limitations warrant consideration. First, while our use of the TwinLife dataset provides a comprehensive view, it is limited to Germany, and the findings may not generalize to other cultural or socioeconomic contexts. Second, our analysis is cross-sectional, which limits the ability to explore the dynamics of GxE interactions over time. Future research should explore the mechanisms underlying the observed GxE interactions in greater detail. Investigating how specific neighborhood factors, such as social capital, collective efficacy, and institutional resources, interact with genetic predispositions could provide deeper insights. Additionally, extending this research to diverse populations and employing longitudinal designs would enhance our understanding of the complex interplay between genetics and environment in shaping life satisfaction.

## 6. Conclusion

This study contributes to the growing literature on the determinants of life satisfaction by highlighting the significant roles of both genetic predispositions and neighborhood conditions. Our findings underscore the importance of considering gene-environment interactions when studying life satisfaction and suggest that the effects of genetic endowments can vary significantly depending on the environmental context. By advancing our understanding of these interactions, we can better inform policies and interventions aimed at enhancing life satisfaction and overall well-being.

## Acknowledgements

This work was made possible by the participants of the German Twin Family Panel (TwinLife). The TwinLife study was funded by the German Research Foundation (Grant No. 220286500), with funding awarded to Martin Diewald, Christian Kandler, Frank M. Spinath, Bastian Mönkediek, and Rainer Riemann. The molecular genetic extension of TwinLife was also funded by the German Research Foundation (Grant No. 428902522), with funding awarded to Martin Diewald, Peter Krawitz, Markus M. Nöthen, Rainer Riemann, and Frank M. Spinath. The funders had no role in the study design, data collection and analysis, decision to publish, or manuscript preparation. YL and MR are supported by the TwinLife project, while NVH is funded bythe European Union’s HORIZON-MSCA-2021-DN-01 programme (Grant No. 101073237). We would like to thank Charlotte Pahnke, Andreas Forstner, Markus Nöthen, Carlo Maj, and Shirin Zare for their contributions to DNA extraction and polygenic score calculations. We also extend our appreciation to Anita Kottwitz from the TwinLife data management team for her invaluable support.

Additionally, we are grateful for the insightful feedback provided by colleagues during the TwinLife Epigenetic Change Satellite (TECS) and TwinSNPs organizational meetings. NVH is now affiliated with the Max Planck Research Group Biosocial – Biology, Social Disparities, and Development; Max Planck Institute for Human Development; Lentzeallee 94, 14195 Berlin, Germany.

